# An Experimental Analysis of the Fine-Scale Effects of Nest Ectoparasites on Incubation Behavior

**DOI:** 10.1101/2022.07.09.499424

**Authors:** Amanda K. Hund, Kelley A. McCahill, Mara Hernandez, Sheela P. Turbek, Daniel R. Ardia, Ryan C. Terrien, Rebecca J. Safran

**Affiliations:** University of Colorado, Department of Ecology and Evolutionary Biology, Boulder, Colorado, USA; Franklin and Marshall College, Department of Biology, Lancaster, Pennsylvania, USA; Carleton College, Department of Physics and Astronomy, Northfield, Minnesota, USA; University of Minnesota, Department of Ecology, Evolution and Behavior, St Paul, Minnesota, USA; National Ecological Observatory Network, Boulder, Colorado, USA; Avon Wildlife Trust, Bristol, England, UK; Colorado State University, Department of Biology, Fort Collins, CO, USA

**Keywords:** Incubation, Parental Care, Parasites, Ectoparasites, Nest Mites, Barn Swallow

## Abstract

Avian incubation is a highly complex, adjustable behavior essential to embryo development and survival. When incubating, parents face a tradeoff between investing in incubation to maintain optimal temperatures for egg development or in self-maintenance behaviors to ensure their own survival and future reproduction. Because nest ectoparasites are costly and can reduce nestling quality and survival, infections could shift parental investment in current vs. future offspring. However, it is not well understood whether birds change investment in incubation in response to nest ectoparasitism, particularly in the context of other factors that are known to influence incubation behavior, such as ambient temperature, clutch size, and embryo development. We hypothesized that parents could respond to nest parasites by 1) investing more in incubation to promote the development of high-quality offspring to help offset the cost of parasites, 2) investing less in incubation or even abandoning their nest during incubation to save resources for future reproduction, or 3) being unresponsive to parasite infections, as incubation is more constrained by other factors. We tested these hypotheses by experimentally removing and adding mites in barn swallow nests at the start of incubation and deploying thermocouple eggs to measure egg temperatures at one-minute intervals until hatching. We found that while ambient temperature, clutch initiation date, embryo age, and clutch size were the main factors driving variation in egg conditions and parental incubation behavior, parasitized nests had higher mean egg temperatures, which could aid in nestling immune development. However, despite more optimal developmental temperatures, eggs in parasitized nests also had lower hatching success. Our results indicate that incubation is a dynamic behavior that is largely driven by the constraints of maintaining appropriate egg temperatures for development. Although quite costly upon hatching, ectoparasites appear to play a relatively minor role in driving variation in parental incubation investment.

**LAY SUMMARY:**

- Incubation is a complex behavior shaped by both internal and external factors.
- Ectoparasites often decrease quality and survival of nestlings and could influence investment in incubation because of tradeoffs in how parents spend energy.
- We manipulated parasites in barn swallow nests and used fake eggs with temperature sensors to collect data on egg temperatures and parental behavior throughout incubation.
- We found that other factors were the main drivers of variation in incubation behavior, but that eggs in parasite nests were warmer on average, which could help nestlings better cope with parasites upon hatching.
- Despite more optimal developmental temperatures, eggs in parasitized nests had lower hatching success.
- Although quite costly to nestlings, ectoparasites played a minor role in shaping parental incubation behavior.

## INTRODUCTION

Avian incubation is a complex and dynamic behavior that requires continuous regulation by the incubating parent to ensure optimal egg temperatures for embryo development (Cooper & Voss 2013). However, investment in incubation is also shaped by important tradeoffs driven by energetic constraints (Bryan and Bryant 1999; Reid *et al*. 2000; Pérez *et al*. 2008; Ardia *et al*. 2009; Cooper and Voss 2013). From an energetic perspective, incubating parents alternate between energy-intensive on-bouts that warm eggs and maintain optimal temperatures for development, and off-bouts which allow for self-maintenance behavior, including foraging and preening (Álvarez and Barba 2014; Ardia *et al*. 2009; Cooper and Voss 2013; MacDonald *et al*. 2012). Studies show that within species higher mean egg temperatures directly correlate with faster embryonic development, shorter incubation periods (days to hatch) and a corresponding reduction in egg predation, and increased survival and quality of nestlings (Martin *et al*. 2007; DuRant *et al*. 2013; Hepp *et al*. 2015; Wada *et al*. 2015). At the same time, frequent trips on and off the nest are energetically costly especially due to the energy cost of rewarming eggs, depleting the body condition and energy reserves of incubating parents, and increase predation risk (Visser & Lessells 2001; Hanssen *et al*. 2005; Nord & Nilsson 2012; Nord & Williams 2015).

Under life history theory, parents should adopt an incubation strategy that balances embryonic development while maximizing survival and lifetime fitness (Voss *et al*. 2006; Ardia *et al*. 2010). Following this, parents may either invest heavily in incubation, using limited resources and risking mortality to maximize current reproductive success, or invest less in incubation and reallocate energy towards survival and future reproductive attempts (Reid *et al*. 2000; Martin 2002). Many external and internal factors can shift the costs and benefits associated with incubation investment and alter how parents invest in incubation in real time. These factors include ambient temperature (Conway & Martin 2000; Voss *et al*. 2006; Ardia *et al*. 2009; Amininasab *et al*. 2016), timing within the breeding season (Ardia *et al*. 2006; Cooper & Voss 2013), parental age or experience (Ardia & Clotfelter 2007; Bogdanova *et al*. 2007; Williams *et al*. 2020), parental condition (Wiebe *et al*. 1998; Ardia *et al*. 2006; Voss *et al*. 2006), clutch size (Engstrand & Bryant 2002; Reid *et al*. 2002; Dobbs *et al*. 2006), and embryo age (Boulton & Cassey 2012; Cooper & Voss 2013). However, a potentially important factor that has been largely overlooked in our understanding of parental investment in incubation is the presence of nest ectoparasites.

Nest ectoparasites are ubiquitous in wild bird populations and often have important negative impacts on nestling development, short and long-term survival, body condition, immunity, and overall quality (Lehmann 1993; Nilsson 2003; Fitze *et al*. 2004; Owen *et al*. 2010; Brommer *et al*. 2011). In many instances, these parasites can also be costly to the parents themselves by taking blood meals from the adults, stimulating an immune response, and reducing investment in other behaviors by increasing preening behaviors (Møller 1993; De Coster *et al*. 2010; Cantarero *et al*. 2013; López-Arrabé *et al*. 2015). In addition, nest parasites can influence other parental care behaviors, including provisioning, nest sanitation, and preening during the nestling period (Christe *et al*. 1996a; Hund *et al*. 2015a; Tripet and Richner 1997). However, it is less clear whether parents dynamically respond to the presence of ectoparasites by changing their incubation behavior during the egg phase of reproduction.

Relatively few studies have examined how nest parasites directly impact incubation behavior (Table 1). In some species, parents appear to shift investment away from incubation in response to nest infection, thereby slowing embryonic development and prolonging incubation periods. In other species, however, parents are unresponsive to nest infections, suggesting that other factors are more important in driving variation in incubation investment (Table 1). While results from past studies have been mixed, we are still missing important details regarding parental responses to nest parasitism during incubation. First, the majority of studies assess incubation investment by either measuring total incubation period (days to hatch) or parental behavior for a few hours at one time point during incubation. These coarse indicators cannot capture the full range of variation in incubation investment, which can depend on changing environmental conditions and subtle dynamics of embryonic development (Londoño *et al*. 2008; Morrison *et al*. 2009). Given this, the detailed measurement of egg temperatures and incubation rhythms across the incubation period would allow for a finer scale assessment of how parents respond to parasites. Second, it is well understood that parents adjust incubation behavior based on a suite of internal and external factors, including body condition, ambient temperature, clutch size, embryo age, and time in the breeding season (Conway & Martin 2000; Deeming 2002; Engstrand & Bryant 2002; Nord & Williams 2015). Given that these factors likely explain a large portion of variation in incubation behavior, assessing the effect of parasites relative to these other factors would allow us to more clearly test whether parents are responding to infections and to what degree.

**Table 1.** Previous studies examining the effect of nest ectoparasites on parental incubation behavior. The table includes the avian and parasite species studied, how incubation behavior was quantified, whether the effect of parasites was investigated relative to other factors that likely influence incubation behavior, and how parasites influenced incubation measures. All of these studies manipulated parasites in some way, either by comparing disinfected nests to natural nests or doing controlled infections of parasites

| Avian Species | Nest Ectoparasite(s) | Measure(s) of Incubation | Relative to other factors? | Effect of parasites on incubation | Citation |
| --- | --- | --- | --- | --- | --- |
| Arctic barnacle goose (*Branta leucopsis*) | Fleas (*Ceratophyllus vagabundus vagabundus*) | Daily average nest temp & temp SD (ibuttons), Daily number of off-bouts & total time off nest (wildlife camera, 5 min minimum interval) | None | No difference in average temp or temp SD, off-bout length or total time away from nest. | de Jong et al. 2019 |
| Barn swallow, European (*Hirundo rustica rustica*) | Tropical fowl mites (*Ornithonyssus bursa*) | Length of incubation period (days) | None | Prolonged incubation period | Møller 1990 |
|  |  | Length of incubation period (days) | Clutch Size | Prolonged incubation period | Møller 1993 |
| Blue Tits (*Cyanistes caeruleus*) | Hen Flea (*Ceratophyllus gallinae*) | Total amount of time attending eggs (camera, 3 hours on day 6 of incubation) | Clutch Size, Male feeding frequency | No difference in total amount of time attending eggs | Tripet et al. 2002 |
| Great Tit (*Parus major*) | Hen Flea (*Ceratophyllus gallinae*) | Length of incubation period (days) | None | No difference in incubation period | Dufva and Allander 1996 |
|  |  | Length of incubation period (days) | None | Prolonged incubation period | Fitze et al. 2004 |
| Pied flycatcher (*Ficedula hypoleuca*) | Natural communities (mites, blowflies, hen fleas) | Total time on the nest, mean length on/off-bouts (camera, 90 min on day 7 of incubation) | None | No difference in total time on the nest or mean length of on/off-bouts | Cantarero et al. 2013 |
| Tree Swallow (*Tachycineta bicolor*) | Bird flea (*Ceratophyllus idius*) | Length of incubation period (days) | None | No difference in incubation period | Rendell and Verbeek 1996 |

In this study, we measured these factors while experimentally manipulating parasites and tracking fine-scale egg temperatures and incubation rhythms over time to fill a key gap in our understanding of how parasites shape investment in incubation. Specifically, we experimentally manipulated blood-feeding mites in the nests of North American barn swallows (*Hirundo rustica erythrogaster*) at the start of the incubation period and used fake eggs with an internal thermocouple probe to monitor egg temperatures at one-minute intervals and quantify daily incubation rhythms.

We had three hypotheses regarding how ectoparasites might tip the balance of life history tradeoffs and change parental investment in incubation: 1) Given that nest parasites lower the quality and survival of nestlings, parents may invest less in incubation or even abandon nests following infection, prioritizing self-maintenance to save zxresources for future reproduction. 2) Parents may limit the cost of ectoparasites to nestlings by investing more in incubation to produce more robust offspring (compensation). While compensation in the face of nest ectoparasites has not yet been shown for incubation behavior, it has been found in some species for nestling provisioning (Tripet *et al*. 1997, 2002; Bouslama *et al*. 2002; Hund *et al*. 2015a). 3). Finally, parents may not change incubation behavior in response to parasites. Given the developmental constraints of eggs (which already represent a substantial investment of resources), the relative impact of parasites may be small compared to other factors (i.e. ambient temperature) that dictate incubation behavior in short lived passerines with limited reproductive opportunities.

## METHODS

### Study System

The North American barn swallow is a widespread semi-colonial migratory passerine that build mud-cup nests exclusively in association with human structures, such as barns and bridges. Breeding pairs have one to three breeding attempts per year, laying four to five eggs on average per clutch (Brown & Brown 1999), although clutch sizes as large as six or seven do occur (*author, pers. obs.)*. While females perform the majority of incubation and are the only sex that develops brood patches, barn swallows have intermittent biparental incubation (Smith & Montgomerie 1992; Turbek *et al*. 2019). Females typically lay one egg per day and begin incubating after laying the penultimate egg (Banbura & Zielinski 1995). The incubation period ranges from 12-17 days and both males and females provision and care for altricial nestlings upon hatching.

Our study population of barn swallows is parasitized by the blood feeding northern fowl mite (*Ornithonyssus sylviarum).* These mites feed primarily on nestlings, where they are readily seen taking blood meals, but are rarely found on adults, including incubating females captured directly off the nest (Hund *et al*. 2021). This suggests that the mites have little direct physiological or immunological impact on adult birds (caused by feeding). These mites complete their entire life cycle in the nest, overwintering in nests between breeding seasons (Hund *et al*. 2015a, 2021). Mite populations in nests can proliferate rapidly when blood meals are available; the entire life cycle from egg to reproductive adult takes five to twelve days and female mites lay multiple clutches of two to five eggs with a female-biased sex ratio (∼5:1) (Richner & Heeb 1995; Proctor & Owens 2000; Szabó *et al*. 2002; Mcculloch & Owen 2012). Additionally, these mites have haplodiploid sex determination, such that virgin females can lay unfertilized eggs, which result in male offspring that they can then mate with to begin an infestation (Mcculloch & Owen 2012). Thus, even at low initial infestation levels, the mite population in a nest can change dramatically across the developmental period of barn swallow nestlings (Dube *et al*. 2018).

### Field Methods

This study took place at 24 breeding colonies in Boulder County, Colorado during the summer of 2016 (May - July). We captured adult birds using mist nests and targeted night captures and gave each bird a numbered aluminum leg band (United States Geological Survey) and a unique combination of color bands that allowed for individual identification. We measured mass and wing length for each adult. We were unable to capture all females at a consistent time relative to the start of incubation (some were captured up to 29 days after), thus we decided not to include adult body condition in our analyses.

By observing color bands, we identified social pairs and assigned them to nests. We checked all nests at study colonies every three days to determine when pairs began lining their nests with feathers and then monitored active nests daily to determine clutch initiation and completion dates. During incubation, we checked nests every four to five days to minimize disturbance while still monitoring for predation events. At the end of incubation (12 days after clutch completion) we checked nests daily to determine hatch date. Nest fate was marked as “hatched” if at least one egg hatched, “predated” if eggs disappeared during the incubation period (presumably eaten), and “abandoned” if eggs remained in the nest but failed to hatch (after 18 days), were cold, and observations indicated that parental nest attendance had stopped.

The nests used in this study were part of a larger egg cross-fostering experiment studying how the developmental environment influences color expression in nestlings. Briefly, we paired experimental nests by clutch completion date, and on the morning of the third day of incubation, the smallest and largest eggs were exchanged across paired nests. Because eggs were reciprocally exchanged between nests, clutch size remained the same. We also included control nests, where eggs were handled in the same way but not exchanged. This allows us to confirm that the cross-foster treatment did not influence parental incubation behavior. As part of this study, we manipulated mites in nests to create parasitized and disinfected treatments (n=130 nests, 58 parasitized, 71 disinfected, 13 of which were control nests). All control nests received the disinfected treatment to isolate any effects of cross-fostering from the impacts of parasites.

For a subset of experimental nests, we installed temperature data loggers to track incubation behavior (n=96 nests, 43 parasitized and 53 disinfected, 12 of which were control nests). Nests without data loggers had the same fake egg tied into place, such that clutch size was consistently increased by one across all nests in the experiment. We checked nests to determine hatching success when nestlings were three days old. While our data come from an egg reciprocal cross-foster study, this manuscript focuses on the parasite manipulation and measures of incubation behavior and hatching success.

### Parasite Manipulations

On the third day of incubation, eggs were removed and all nests were disinfected by heating them and the surrounding substrate to 120°C with an industrial heat gun to kill any naturally occurring parasites (Hund *et al*. 2015b). While nests cooled, we placed fake plastic eggs in the nest so returning parents would not observe that eggs were missing. We returned the real eggs to the nest after it had cooled (∼5min). We then randomly assigned one nest in each cross-foster pair to a parasite treatment, where 100 live field-collected mites were added into the nest cup. The mites used in this experiment were collected from non-experimental nests, vacuumed into tubes, and stored in a humidified refrigerator (1.6°C) for up to seven days until they were added into nests (Hund *et al*. 2015a, 2021).

### Temperature data loggers

To monitor egg temperatures and incubation behavior, we deployed plastic model eggs with internal thermocouple temperature data loggers that recorded temperatures at one-minute intervals (OM-EL-USB-TC, Omega Engineering, Inc., hereafter referred to as omegga loggers). The egg part of each omegga logger was painted to look like a barn swallow egg and was equivalent in both size and shape (Figure 1). These fake eggs were filled with a gel (Clear Glide wire pulling lubricant) that closely mimics the thermal properties of albumin in a real egg (Ardia *et al*. 2010). To prevent females from rejecting or attempting to turn or center these model eggs, we tied them into place by passing a thread attached to the bottom of the fake egg carefully through the nest with a needle and securing it directly below. The data logger was placed in a plastic bag and secured near the nest (∼30-60cm away). Birds quickly acclimated to the presence of the fake egg and logger and begin incubating again shortly after omegga installation. Omegga loggers remained in the nest until three days post hatching. Similar thermocouple eggs have been used in tree swallows (*Tachycineta bicolor)* to study incubation, and it was found that the addition of an artificial egg did not significantly impact female incubation behavior or nest attendance (Ardia *et al*. 2009, 2010).

**Figure 1.**
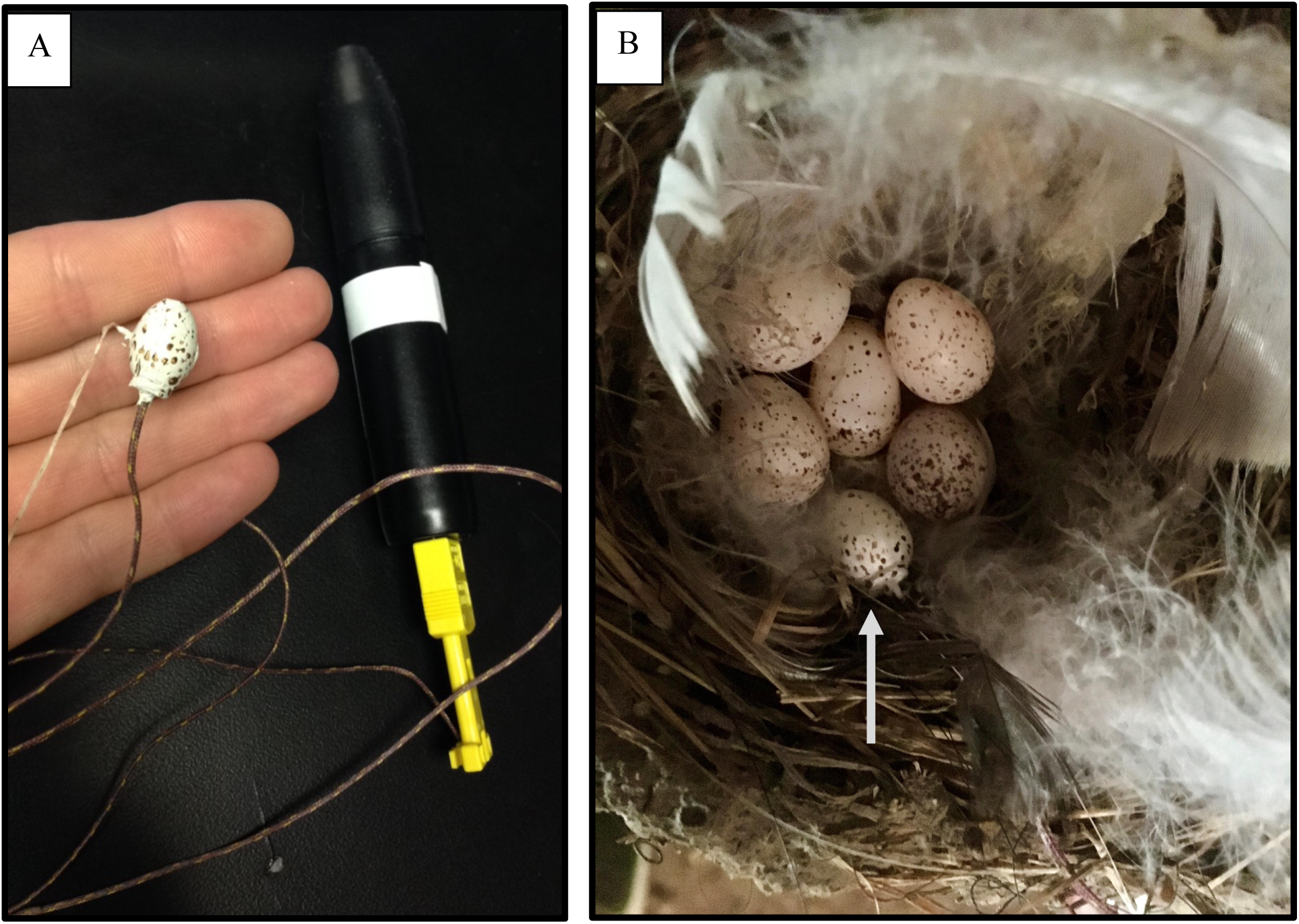
A) Picture of an omegga data logger showing the fake egg with internal thermocouple probe, thread for securing the egg, and thermocouple wire attached to a data logger. B) Omegga logger installed in a barn swallow nest. The arrow indicates the fake egg.

### Processing omegga data

To ensure that incubation patterns extracted from egg temperature data reflected real parental behavior, we observed and recorded incubation behavior for 16 nests that had omegga loggers installed. After a 15min habituation period, we observed nests for 1 hour and recorded the movements of parents to and from the nest. When we compared these field observations to the corresponding temperature graphs collected from our loggers, we found that there was a temperature detection delay of about two minutes (i.e. egg temperature decreasing after the parent left the nest, or increasing once the parent returned to the nest), and that only changes >0.5°C were reliable indications of parents leaving or returning to nests. This was consistent across all 16 nests and allowed for a stringent set of editing rules to ensure that we accurately extracted behavioral data from our omegga temperature readings.

We analyzed incubation behavior for nests that successfully hatched and for which the omegga loggers worked and remained in place (n=68, 33 parasitized, 35 disinfected nests, 9 of which were controls). As incubating females typically spend the night on their nests, most variation in incubation behavior occurs during the day. Given this, we split our temperature data into day and night categories using sunrise and sunset times for each date (found using the Astroplan software package, (Morris *et al*. 2018)). We analyzed egg temperature data for both day and night but used only the day data to assess behavioral rhythms (trips on and off the nest). To allow for habituation to the omegga logger and a return to normal behavior after disturbance generated by working near the nest, we excluded data collected on the day we installed the loggers. We also excluded any readings collected on or after hatch day, as we were interested specifically in incubation behavior.

For the egg-temperature data (extracted from our omegga loggers), we first examined the mean egg temperature for each day and night. We then categorized each one-minute reading into one of four biologically relevant temperature zones (Conway & Martin 1999; Cones *et al*. 2020): 1) <26°C, at which embryos arrest development (referred to as the “cold” zone), 2) 26-34°C, at which slow and sometimes detrimental embryonic development occurs (referred to as the “lukewarm” zone), 3) 34.1-40.5 °C, which is the optimal temperature range for embryonic development in most birds, though this can vary some by species (referred to as the “warm” zone), and 4) >40.5, at which temperatures are too hot and can be determinantal or even lethal to embryos (referred to as the “hot” zone). Once all reads were categorized, we counted the number of readings (or minutes) that eggs spent in each of these different zones for each day and night of incubation.

To extract incubation behavior from our temperature data, we first used Rhythm (1.0) to detect egg cooling and re-warming periods (Cooper *et al*. 2005). We used the following detector settings based on our behavioral observations and past studies that have used similar loggers: a minimum off-bout duration of two minutes, minimum depth of one degree, cooling and re-warming period slopes of one-quarter degree per minute, and an automatic timeout setting of ten minutes (Ardia *et al*. 2009; Cooper *et al*. 2005b; Cooper and Voss 2013). We then used Raven Pro (64 1.5) to visualize daily temperature graphs with rewarming, equilibrium, and cooling periods. Daily temperature graphs were manually checked and sometimes edited to ensure period selections were correct. We averaged extracted incubation behaviors for each day, leading to repeated daily measures within each nest.

### iButtons

We deployed iButtons (Maxim Integrated) at each site that recorded ambient temperatures at 30min intervals throughout the breeding season. We hung iButtons in small mesh bags in different parts of each structure to capture the particular microclimate that nests were experiencing (e.g. in rafters further or closer to the apex of a roof; at the entrance vs. middle of a culvert or bridge, 2-10 iButtons per site depending on the size and shape of the structure). We then matched each nest to the nearest iButton to pair our incubation data with ambient temperature data. We separated iButton data by daily sunrise and sunset time to align with our incubation measures.

### Unhatched eggs

We collected intact unhatched eggs that remained in experimental nests when nestlings were nine days old (n=28 eggs from parasitized nests and 42 eggs from disinfected nests) and stored them in the freezer. Unhatched eggs were often expelled from nests or found broken within the nest; Thus, the unhatched eggs that we collected were a subset of all the eggs that failed to hatch from our experimental nests. We dissected these eggs to determine the developmental stage at which the embryos died. We divided embryonic development into six different stages that were easily distinguishable using key features, from unfertilized egg (1) to ready to hatch (6) (See figure S1 for a chart depicting the six stages). To create this scale, we used published developmental charts from other passerines and dissected frozen known-age barn swallow eggs from a previous experiment where eggs were collected during incubation (Levin *et al*. 2018). The same person (MH) dissected all eggs while blind to treatment.

### Statistical Analysis

#### Reciprocal cross-foster design

To confirm that switching eggs between nests did not influence incubation behavior, we compared measurements of egg temperatures and incubation rhythms between control and cross-fostered nests within the disinfected treatment (n=23 cross-fostered and 12 control nests). We used linear mixed models (LMM) and general linear mixed models (GLMM) to compare: daily and nightly mean egg temperatures, mean number of off-bouts, mean off-bout length, proportion of time parents spent on the nest, days to hatch (length of incubation period), and the per-egg probability of hatching failure (unhatched eggs / clutch size). To account for repeated measures within a nest, we included embryo age nested within nestID as random effects. To account for nests sharing the same breeding site, we also included site as a random effect. Because some of our control nests were laid slightly later in the season, we included clutch initiation date (Julian Date, or days since Jan 1) in these models.

#### Nest Abandonment

To assess whether the parasite treatment caused parents to abandon nests at higher rates during the incubation period, we used the full data set (n=130 nests, 58 parasitized, 71 disinfected), but excluded nests that were eaten by predators during incubation (7 parasitized and 8 disinfected nests). We carried out a GLMM (binomial), with whether or not a nest was abandoned as the response variable, treatment as a fixed effect, and site as a random effect.

#### Egg Temperature

Using our raw data from the omegga loggers (n=68, 33 parasitized, 35 disinfected nests), we first wanted to see if parasite treatment influenced egg temperature and the developmental conditions that embryos experienced. We compared daily and nightly mean egg temperatures as our response variables using LMMs. Mean night temperature was square root transformed to improve model stability. We also compared the time-spent (count of one-minute reads) in each of our four temperature zones using GLMMs with a Poisson distribution with a log link. Several of these models were zero-inflated and overdispersed; We thus included an individual level random effect. We included the following fixed effects: parasite treatment, clutch size, clutch initiation date, embryo age (days since the start of incubation) and mean ambient temperature. For several of our temperature zone models, we were unable to include all these fixed effects because the models would not converge. Given this, we removed clutch size, which data exploration indicated had little effect on time spent in different temperature zones. All our models included embryo age nested within nestID (to account for repeated measures within a nest) and site as random effects.

#### Incubation Behavior

We next tested if parasites influenced our measures of incubation behavior (trips on and off the nest). We used the daily average number of off-bouts (LMM), the average off-bout length (LMM, response log transformed), and the total daily proportion of time parents spent on the nest as our response variables (GLMM, beta with a “logit” link, (Douma & Weedon 2019)). We included the following fixed effects: parasite treatment, clutch size, clutch initiation date, embryo age, and mean ambient temperature. We again included embryo age nested within nestID and site as random effects.

#### Outcomes of Incubation

In addition to measures of egg temperature and parental behavior, we tested whether parasites influenced the outcomes of incubation. To do this, we looked at the egg-level probability of hatching failure (GLMM, binomial with a “logit” link) and the number of days eggs took to hatch (length of the incubation period, GLMM, Poisson), as less attentive incubation behavior can slow development time, which often results in lower quality offspring (Gorman *et al*. 2005; Olson *et al*. 2006). Many of our different measures of egg temperature and behavior were correlated. Given this, we include the following fixed effects to assess the relative role of parasites: parasite treatment, daily mean egg temperature, clutch size, clutch initiation date, and average number of off-bouts. In our model examining hatching failure, we were unable to include all of these fixed effects because the model would not converge. Given this, we removed clutch initiation date as a fixed effect, which data exploration indicated had little effect on the probability of hatching failure. We again included embryo age nested within nestID and site as random effects.

#### Unhatched eggs

We compared the embryonic stage at which embryos died in unhatched eggs between treatment groups using a cumulative link mixed model for ordinal data, from the package “Ordinal” (Christensen 2019). We included site as a random effect in this model. There were cases where we collected more than one unhatched egg from the same nest. As the Ordinal package does not support nested random effects, we reran our model as a LMM with nest and site as nested random effects. These models produced very similar results.

#### Modeling strategy

For each of the models described above, we first checked for correlations between fixed effects to ensure they could be included in the same model and then centered and scaled all numerical fixed effects to have a mean of zero and a standard deviation of one, which allows us to directly compare parameter estimates within a model (Zuur *et al*. 2009). For each model, we checked for issues like overdispersion, examined residuals for appropriate fit, and confirmed other necessary assumptions. Because we were specifically interested in testing the role of parasite treatment relative to the other fixed effects, we built a full model for each response variable and then a second model where parasite treatment was removed. We then compared these models with a log-likelihood test and Akaike Information Criterion (AIC) values. In cases where the inclusion of parasite treatment improved the model, we assessed its significance and relative effect by comparing the parameter estimates between different fixed effects. We report significant results in the main text, and full model results can be found in the supplement. When possible, we also report the conditional r-squared value (R^2^c, associated with the fixed effects plus the random effects). All analyses were done in R, version 4.0.2 (R core Team 2020), with the MuMin (Bartroń 2017), lme4 (Bates et al. 2015), and glmmTMB (Brooks *et al*. 2017) packages.

## RESULTS

### Reciprocal cross-foster design

None of our measures of egg temperature, incubation behavior, or outcomes of incubation differed between control and cross-fostered nests within the disinfected treatment (Table S1). This suggests that exchanging eggs between nests did not significantly influence incubation behavior. Given this, we included the control nests as part of the disinfected treatment for all analyses below.

### Nest Abandonment

In the parasitized treatment, 19.2% of nests were abandoned before eggs hatched compared to 12.7% of nests in the disinfected treatment, though this difference was not statistically significant (β = 0.21, SE = 0.62, *F* = 0.11, *P* = 0.74, R^2^c = 0.29).

### Egg Temperature

Daily mean egg temperature was significantly associated with parasite treatment; eggs in parasitized nests were warmer on average than eggs in disinfected nests (β= 0.85, SE = 0.43, *F* = 3.80, *P* = 0.05, R^2^c = 0.95; LSM: parasitized= 34.6°C ± 0.37, disinfected= 33.8°C ± 0.38). Including parasite treatment as a fixed effect improved model fit (ΔAIC = 1.8, *X*^2^ = 3.8, *P* = 0.05). Daily ambient temperature had the largest effect on egg temperature (Figure 2); intuitively, eggs were warmer when ambient temperatures were warmer (β = 1.19, SE = 0.04, *F* = 941.75, *P* < 0.001, R^2^c = 0.95). Clutch initiation date was also important, as eggs laid later in the season were warmer (β= 0.75, SE = 0.23, *F* = 11.01, *P* = 0.001, R^2^c = 0.95). We note that, while ambient temperatures do tend to be warmer later in the breeding season, these fixed effects were not tightly correlated (R^2^= 0.22), suggesting that increased egg temperature in later nests was not due solely to increasing seasonal temperatures. Nightly mean egg temperature was also warmer in parasitized nests (LSM: parasitized= 32.9°C ± 0.62, disinfected= 31.6°C ± 0.64), but this effect was not statistically significant (β = −0.25, SE = 0.15, *F* = 2.93, *P* =0.09, R^2^c = 0.74) and including treatment did not improve the model fit. Similar to our findings during the day, nightly egg temperatures were largely driven by ambient temperature (Figure S2, β= −0.13, SE = 0.03, *F* =20.97, *P* <0.001, R^2^c =0.74) and clutch initiation date (β= −0.20, SE = 0.08, *F* =6.11, *P* =0.02, R^2^c =0.74).

**Figure 2.**
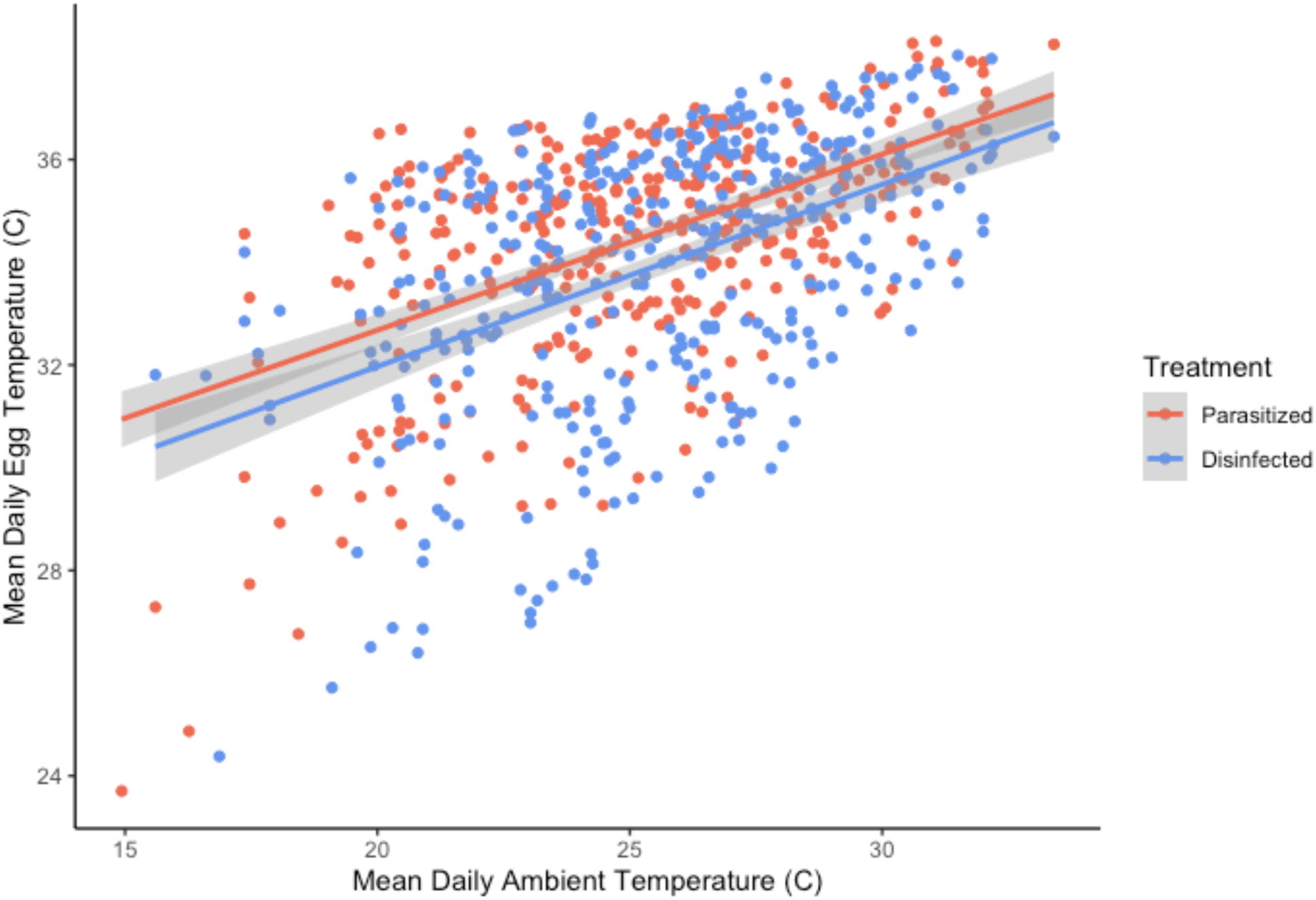
Relationship between daytime mean egg temperature, measured with omegga loggers, and ambient temperature near the nest, measured with iButtons, separated by parasite treatment group. Parasitized nests were, on average, warmer than disinfected nests; however, egg temperature was largely driven by ambient temperature. Figure and trend lines were made with raw data. Analyses presented in the text include additional fixed effects and embryo age (nested within nest) and site as random effects to account for repeated measures within the same nest and multiple nests from same breeding colonies.

Ambient temperature was the dominant factor predicting time spent in all temperature zones, during both the day and night (Figure S3). As expected, eggs spent less time in the cold and lukewarm zones and more time in the warm and hot zones as ambient temperature got warmer (see Table S2 for full model results). Treatment did influence the amount of time eggs spent in the warm zone at night, with parasitized nests spending more time on average in this ideal zone compared to disinfected nests (β= 2.08, SE = 0.82, *F* =2.93, *P* =0.01, disinfected mean: 224 mins, parasitized mean: 258 mins), and including parasite treatment as a fixed effect improved the fit of this model (ΔAIC = 3.5, *X*^2^ = 5.5, *P* = 0.02). Time spent in the warm zone at night was also influenced by clutch initiation date, where time increased as eggs were laid later in the season (β= −1.41, SE = 0.45, *F* =15.19, *P* =0.001), and embryo age, where eggs spent less time in this zone as embryos aged (β= −0.23, SE = 0.07, *F* =0.06, *P* =0.002). The time that eggs spent in the cold temperature zone during the day was also significantly predicted by both clutch initiation date (β = −1.05, SE = 0.49, *F* =11.65, *P* =0.03), and embryo age (β = −0.92, SE = 0.31, *F* =21.46, *P* =0.004); eggs spent less time in the cold zone if they were laid later in the season and as embryos aged.

### Incubation Behavior

Parasite treatment had little effect on measures of daily parental incubation behavior (trips on and off the nest) and all models improved when treatment was removed. The mean number of off-bouts was driven primarily by ambient temperature, where the number of off-bouts decreased with warmer ambient temperatures (β =-5.22, SE = 0.41, *F* = 162.00, *P* < 0.001, R^2^c =0.82). The number of off-bouts also increased as embryos aged (β = 1.45, SE = 0.42, *F* = 11.55, *P* = 0.001, R^2^c =0.82). We found the inverse pattern with mean off-bout length (Figure 3A), which increased with warmer ambient temperatures (β = 123.55, SE = 12.38, *F* = 99.64, *P* < 0.001, R^2^c =0.67). These results suggest that parents took more frequent, shorter, off-bouts when ambient temperatures were cool or embryos were more developed and less frequent, longer, off-bouts when ambient temperatures were warm, or embryos were younger. Similarly, parents spent a greater proportion of time on the nest when ambient temperatures were cooler (Figure 3B, β = −0.06, SE = 0.01, *Z* =, *P* < 0.001).

**Figure 3.**
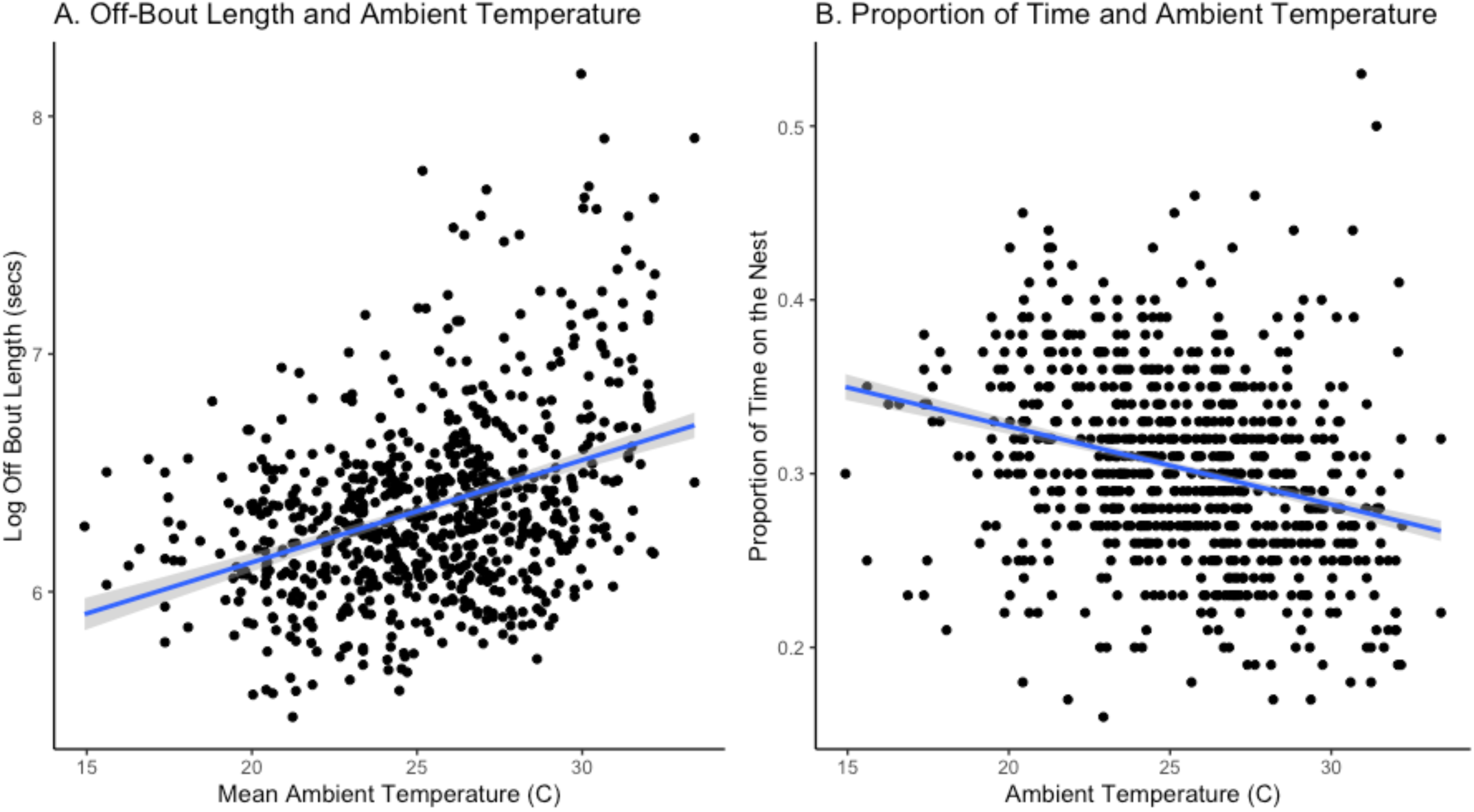
Relationship between ambient temperature and two measures of parental incubation behavior: A) average daily off-bout length and B) daily proportion of time on the nest. As temperatures warmed, parents took fewer (not shown), longer off-bouts, and spent a smaller proportion of their time on the nest. Figures and trend lines are made with raw data. Analyses presented in the text include embryo age nested within nest, to account for repeated measures of the same nest, as well as breeding site as random effects.

### Outcomes of Incubation

The probability of egg hatching failure was higher in parasitized nests (β = 0.75, SE = 0.34, *F* = 1.09, *P* = 0.03, R^2^c = 0.27), and including parasite treatment improved model fit (ΔAIC = 2.2, *X*^2^ = 4.20, *P* = 0.04). The probability of hatching failure also increased as clutch size decreased (β = −0.57, SE = 0.14, *F* = 17.00, *P* < 0.001, R^2^c =0.27), increased with the number of off-bouts (β= 0.32, SE = 0.14, *F* = 6.85, *P* = 0.02, R^2^c =0.27), and increased as average daily egg temperature increased (Figure 4, β = 0.71, SE = 0.23, *F* = 13.88, *P* = 0.002, R^2^c = 0.27).

**Figure 4.**
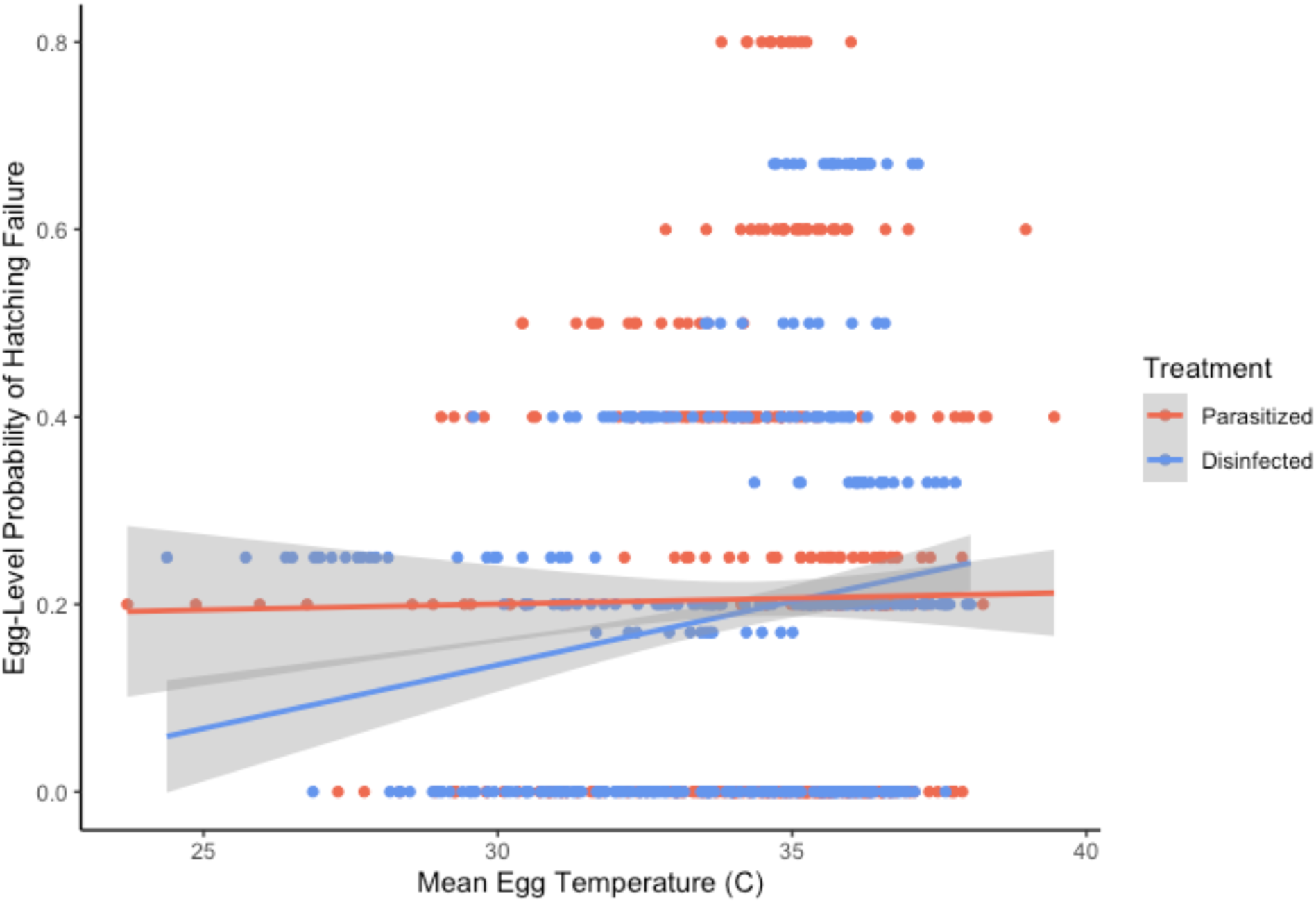
Relationship between the egg-level probability of hatching failure and the mean daily egg temperature, measured with omegga loggers. We highlight the effect of parasite treatment for comparison. At warmer egg temperatures, egg failure rates are similar across treatments, but parasitized nests had higher failure rates at lower mean egg temperatures compared to disinfected nests. Figure and trend lines are made with raw data, but analyses presented in the text include embryo age nested within nest, to account for repeated measures of the same nest, as well as breeding site as random effects.

However, parasite treatment did not have a strong effect on incubation period (days to hatch) and did not improve model fit. The length of the incubation period was largely driven by clutch initiation date; where days to hatch decreased as clutch initiation date moved later into the breeding season (β = −0.03, SE = 0.01, *F* = 6.39, *P* = 0.01).

### Unhatched Eggs

There was no difference between treatment groups in the developmental stage of unhatched eggs (β = −0.26, SE = 0.46, *Z* = −0.57, *P* = 0.57, n=28 eggs from parasitized nests and 42 eggs from disinfected nests). In general, we found that stage one (likely unfertilized) was the most common categorization, and that the second stage (very early development) was the least common (stage one: 29.6%, stage two: 2.9%, stage three: 11.4%, stage four: 18.6%, stage five: 18.6%, stage six: 20.0%).

## DISCUSSION

Nest ectoparasites can have important impacts on nestling condition, short- and long-term survival, and overall quality (Lehmann 1993; Nilsson 2003; Fitze *et al*. 2004; Owen *et al*. 2010; Brommer *et al*. 2011). During the nestling phase, parents are known to shift their behavior in response to parasite infections, including changes in provisioning, nest sanitation, and preening (Christe *et al*. 1996a; Hund *et al*. 2015a; Tripet and Richner 1997). Yet it remains unclear if and how parents respond to nest ectoparasites during incubation. In particular, we lack a detailed view of behavior throughout the incubation period and the relative effect of ectoparasites in the context of other factors that are known to influence incubation behavior, including ambient temperature, clutch size, embryo age, and timing within the breeding season. We had three hypotheses for how parents may respond to the presence of nest ectoparasites during incubation: 1) decrease investment in incubation to save resources for future reproduction, 2) increase investment to offset the cost of ectoparasites to nestlings, or 3) not respond, as other factors are more important in shaping incubation behavior. To test these hypotheses, we experimentally manipulated ectoparasites in the nests of barn swallows early in the incubation period and installed loggers to track egg temperatures at one-minute intervals until hatching.

We found that eggs in parasitized nests were warmer on average and spent more time in the optimal temperature range at night, which could indicate an increased investment in incubation (supporting hypothesis two). However, the relative effect of parasite treatment on egg temperature was small compared to other environmental factors, including ambient temperature and clutch initiation date. We also found that eggs in parasitized nests had lower hatching success, which could support hypothesis one, but this was unrelated to our measures of incubation behavior. This unusual pattern suggests more study is needed to better understand the potential impact of ectoparasites on embryonic development conditions. Hatching success was also associated with ambient temperature and clutch size. Parasite treatment had little effect on measures of incubation rhythms (trips on and off the nest) or incubation period (days to hatch), both of which were largely influenced by factors such as ambient temperature, embryo age, clutch size, and clutch initiation date (supporting hypothesis three). Taken together, it seems that nest ectoparasites did influence some aspects of egg development (egg temperature and hatching success), which could suggest changes in parental investment. However, overall, ectoparasites had a relatively small influence on incubation behavior compared to other biological and environmental factors that are important for egg development.

Warmer egg temperatures in parasitized nests may represent an increased investment by parents. Higher egg temperatures (within the optimal developmental range) have been shown to increase offspring quality, size, body condition, and survival in a number of species (Martin *et al*. 2007; DuRant *et al*. 2013; Hepp *et al*. 2015; Wada *et al*. 2015). Interestingly, warmer egg temperatures have also been associated with increased immune investment and innate immune capacity in nestlings (Ardia *et al*. 2010; Liu *et al*. 2013; Merrill *et al*. 2020). Innate immunity is an important first line of defense against parasites, including blood-feeding ectoparasites (Owen 2010), as it develops rapidly during the nestling period and provides protection before the adaptive branches of the immune system are fully functional (Palacios *et al*. 2009; Stambaugh *et al*. 2011; Killpack *et al*. 2013). It is possible that, by keeping eggs warmer in parasitized nests, parents are preparing nestlings to better deal with ectoparasites upon hatching. There are other examples of maternal effects or parental cues during egg development that provide environmental information and adaptively alter offspring development (Duckworth *et al*. 2015; Mariette & Buchanan 2016). Future studies in this system that couple measures of egg temperature with immune measures in nestlings would provide more insight into whether the temperature differences that we detected are shifting offspring phenotype to adaptively respond to ectoparasites.

The mechanism by which parents are increasing egg temperatures in parasitized nests is currently unclear, as we did not detect differences in our measures of incubation behavior. While trips on and off the nest are a good measure of general investment in incubation, parents may also differ in their behavior while on the nest. Incubating birds can influence egg temperatures by varying their posture and the distance between the eggs and the brood patch (Boulton and Cassey 2012), changing their heart rate and blood flow to the brood patch, and even shivering to warm eggs more quickly (Gabrielsen & Steen 1979; Haftorn *et al*. 1982; Tøien 1993; DuRant *et al*. 2013). Even the size of the brood patch is plastic within individuals (Jónsson et al. 2006, Massaro et al. 2006). For North American barn swallows specifically, we know that males help their mates with incubation (∼9% of total daytime incubation), but do not have brood patches and thus are less efficient at warming eggs (Smith & Montgomerie 1992). Changes in the proportion of time males and females are incubating could shift egg temperatures without changing incubation rhythms. These different behaviors, which we could not quantify with our omegga loggers (besides measuring egg temperatures), could have contributed to differences in egg temperatures across parasite treatments. Future studies that incorporate both temperature loggers and videos of parental behavior could provide additional insight.

Reduced hatching success in parasitized nests, on the other hand, could suggest a decreased investment in incubation, though we detected no differences in our behavioral data. Because our experiment manipulated parasites after eggs were laid, and involved cross-fostering eggs between nests, we can largely exclude maternal effects leading to differences in fertilization, egg size, or composition as contributing to variation in hatching success between treatments. We also found no difference in the age at which embryos died. It is unlikely that warmer egg temperatures in the parasite treatment are related to decreased hatching success. At very hot ambient temperatures, which likely did increase egg mortality, there appears to be little difference between treatment groups (Figure 4), and we found no difference between treatment groups in the amount of time eggs spent in the dangerous “hot” temperature zone. Some studies have also found that nest ectoparasites decrease hatching success (Oppliger et al. 1994, de Lope et al. 1993, Pryor and Castro 2017), though others found no effect, including in European barn swallows (Møller 1990, 1993, Tomás et. al 2006, Sźep and Møller 2000, Heeb et al. 2000, Tripet et al. 2002, Richner et al. 1993, Allander 1998, Dufa and Allander 1996). More recent research suggests that the parasites themselves may influence hatching success by changing the bacterial community and load on the egg surface, leading to increased embryo mortality (Ruiz-Castellano *et al*. 2016; Peralta-Sánchez *et al*. 2018; Tomás *et al*. 2018). Future research into the egg surface microbiome could provide more insight into the link between nest ectoparasites and decreased hatching success in this system.

Because nest parasites decrease hatching success and nestling quality and survival in this system, parents could prioritize self-maintenance and future reproduction by abandoning infected nests altogether. There is evidence of ectoparasites causing nest desertion in other systems at both the egg and nestling phase (Duffey 1983; Oppliger *et al*. 1994; Loye & Carroll 1998; Pryor & Casto 2017). We did find that parasitized nests were abandoned at a higher rate (19.2% compared to 12.7%), but this was not statistically significant. It could be that birds are avoiding parasites when initially selecting nest sites (Mazgajski 2007; Hund *et al*. 2021) but are less likely to abandon their nest if parasite transmission occurs after they have already invested in eggs (as in this experiment). Alternatively, parents could be sensitive to the intensity of the infection. For this study, we added 100 mites to each nest in the parasite treatment, which represents a low to moderate infection level (Hund *et al*. 2021). Because the costs of ectoparasites are typically intensity dependent, there could be a threshold effect, such that parents are more likely to abandon nests at higher infection levels. This could also be true for investment in other aspects of incubation behavior.

Parental incubation behavior in our study was largely driven by ambient temperature, which has been well supported in incubation research (Durant et al. 2013). When ambient temperatures were cool, parents took shorter, more frequent off-bouts and spent a greater proportion of their time on the nest to maintain optimal egg temperatures. As ambient temperatures warmed, parents took longer, less frequent off-bouts and spent less time on the nest overall. We also found that embryo age influenced off-bout length. This is consistent with the idea that both egg cooling rate and thermal tolerance changes as embryos develop. Early in development, heat flow through eggs is determined by conduction, while later in development, blood circulation within the embryo increases heat loss (Turner 1987, Turner 2002). Because birds progress from ectothermy to endothermy as they develop within the egg, the thermal tolerance of embryos also decreases as they approach hatching (Ricklefs and Hainsworth 1968, Webb 1987). Given these constraints, parents shorten off-bouts later in incubation to minimize fluctuation in egg temperatures (Cooper and Voss 2013). Similarly, the length of the incubation period, which has been shown to increase with nest ectoparasites in some studies (including studies done on European barn swallows) (Møller 1990, 1993), did not change with parasite treatment in our study. Instead, the length of the incubation period was determined by both clutch size and clutch initiation date. This is likely driven by slower heat loss from large clutches and warmer ambient temperatures later in the breeding season (Boulton & Cassey 2012).

## Conclusion

We found that ectoparasites influenced aspects of egg development in barn swallows, including warmer egg temperatures in infected nests, which could prepare nestlings to better face ectoparasites upon hatching. However, ectoparasites had little impact on other measures of incubation behavior or development time. Our study suggests that incubation behavior is tightly regulated to respond to environmental conditions and developmental constraints. Despite being quite costly to nestlings, ectoparasites (at moderate infection levels) appear to play a relatively minor role in shaping investment in incubation. By examining the fine-scale effects of nest ectoparasites through time and in the context of other key factors that influence incubation behavior, our experiment enabled us to analyze the relative impact of parasites on parental investment in incubation.

## Data Accessibility

All data and code for this manuscript will be made publicly available on Dryad upon acceptance.

## Author Contributions

AH and RS designed and funded the project, AH, KM, MH, ST carried out the fieldwork, KM characterized incubation rhythms, MH dissected and scored unhatched eggs, DA provided information on how to build and use egg loggers and advised on study design and analysis, RT performed data processing from loggers and ibuttons and helped with coding, AH performed data analysis and made the figures, AH and KM wrote the manuscript, all authors provided feedback on the manuscript.

## Acknowledgements

Thanks to undergraduates W Dube and C Mastrangelo and middle school teachers K Greene, H Reeg, and D Tomlin (funded through NSF DEB-CAREER 1149942, Research Experience for Teachers (RET) to RS), for their help with fieldwork. Special thanks to the site owners for allowing us to work on their farms and to the Snell-Rood lab for providing helpful feedback on this manuscript.

## Ethical Note

This research followed the Animal Behavior Society guidelines for the treatment of animals in research. Observations, captures, and handling of birds was done in accordance with the guidelines set by the University of Colorado Institutional Animal Care and Use Committee (IACUC). IACUC reviewed and approved all methods used in this research (permit number 1303.03). Because Barn Swallows nest in human-made structures where human traffic is common (i.e. barns), they are quite robust to nest disturbances and to handling. Abandonment immediately after nests are checked, eggs are handled, or nestlings are measured is very rare and did not occur with any of the birds in this study. Experimental nest mite manipulations were targeted to be at low or moderate infection levels (typically sublethal to nestlings) relative to naturally occurring infections.

## SUPPLEMENTAL MATERIAL

**Figure S1.**
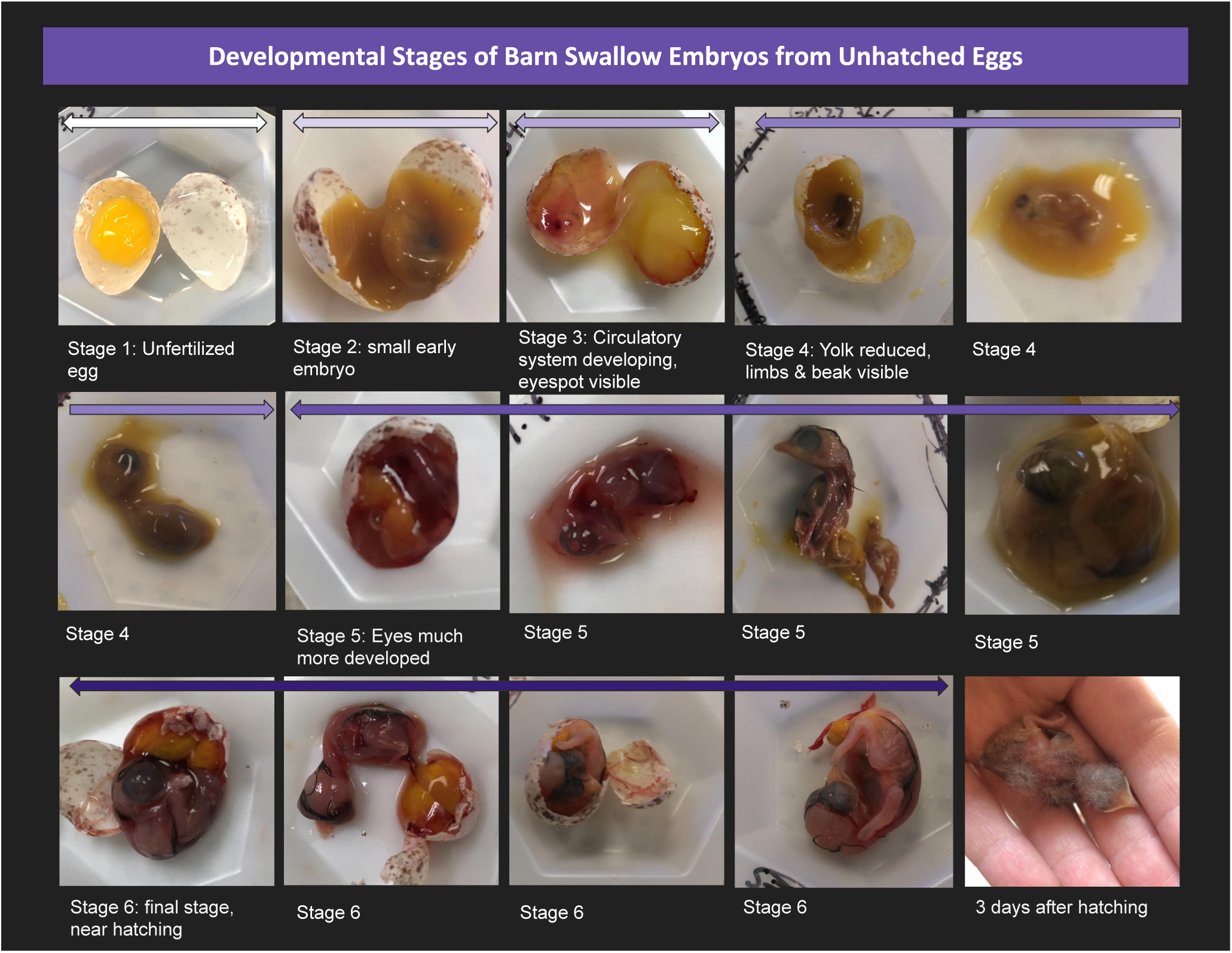
Diagram illustrating barn swallow embryonic development broken into six identifiable stages. Stages were determined by dissecting known-age barn swallow embryos and referencing developmental charts of other passerine species (i.e. zebra finches). We assigned embryos dissected from unhatched eggs collected from our experimental nests to one of these six stages.

### Reciprocal cross-foster design

**Table S1.** Model results comparing control nests to cross-foster nests within the disinfected treatment group. Clutch initiation date was included as a fixed effect because the average clutch initiation date was slightly later in the season for some control nests. All models contained embryo age nested within nest ID and site as random effects.

| Response | Fixed Effect | Estimate | SE | F | P |
| --- | --- | --- | --- | --- | --- |
| Mean daily egg temperature | Cross-fostered | 0.39 | 0.98 | 0.16 | 0.69 |
|  | Clutch Initiation Date | 0.14 | 0.05 | 8.12 | 0.008 |
| Mean nightly egg temperature | Cross-fostered | -0.25 | 0.27 | 0.83 | 0.37 |
|  | Clutch Initiation Date | -0.05 | 0.01 | 13.79 | <0.001 |
| Mean number of off bouts | Cross-fostered | 4.95 | 4.88 | 1.03 | 0.32 |
|  | Clutch Initiation Date | 0.18 | 0.24 | 0.52 | 0.48 |
| Mean off bout length | Cross-fostered | -0.07 | 0.12 | 0.35 | 0.56 |
|  | Clutch Initiation Date | -0.001 | 0.01 | 0.74 | 0.39 |
| Proportion of time on the nest | Cross-fostered | -0.08 | 0.07 | (Z) -0.97 | 0.33 |
|  | Clutch Initiation Date | -0.0005 | 0.003 | (Z) -0.15 | 0.88 |
| Days to hatch | Cross-fostered | -0.02 | 0.03 | 0.43 | 0.57 |
|  | Clutch Initiation Date | -0.003 | 0.002 | 3.44 | 0.07 |
| Probability of hatching failure | Cross-fostered | 30.340 | 31.91 | 0 | 0.34 |
|  | Clutch Initiation Date | 0.11 | 0.02 | 27.02 | <0.001 |

### Egg Temperature

**Table S2.** Full model results for egg temperature and time spent in different temperature zones during the day and night. All models contained embryo age nested within nest ID and site as random effects

| Response | Fixed Effect | Estimate | SE | F | P |
| --- | --- | --- | --- | --- | --- |
| Mean Daily Egg Temperature<br>($R^2c = 0.95$) | Treatment (P) | 0.85 | 0.43 | 3.80 | 0.05 |
|  | Clutch Size | 0.23 | 0.22 | 1.08 | 0.30 |
|  | Clutch Initiation Date | 0.75 | 0.22 | 11.01 | 0.001 |
|  | Embryo Age | -0.01 | 0.05 | 0.08 | 0.77 |
|  | Ambient Temperature | 1.19 | 0.04 | 941.75 | <0.001 |
| Mean Nightly Egg Temperature<br>($R^2c = 0.74$) | Treatment (P) | -0.25 | 0.14 | 2.93 | 0.09 |
|  | Clutch Size | -0.06 | 0.08 | 0.58 | 0.45 |
|  | Clutch Initiation Date | -0.20 | 0.08 | 6.11 | 0.02 |
|  | Embryo Age | 0.003 | 0.02 | 0.02 | 0.88 |
|  | Ambient Temperature | -0.13 | 0.03 | 20.97 | <0.001 |
| Time in Cold (day) | Treatment (P) | -1.05 | 0.94 | 0.19 | 0.27 |
|  | Clutch Initiation Date | -1.05 | 0.49 | 11.65 | 0.03 |
|  | Embryo Age | -0.92 | 0.31 | 21.46 | 0.004 |
|  | Ambient Temperature | -0.84 | 0.21 | 16.69 | <0.001 |
| Time in Lukewarm (day) | Treatment (P) | -0.12 | 0.19 | 0.004 | 0.52 |
|  | Clutch Size | -0.11 | 0.09 | 0.03 | 0.24 |
|  | Clutch Initiation Date | -0.07 | 0.10 | 26.13 | 0.46 |
|  | Embryo Age | 0.008 | 0.03 | 15.15 | 0.73 |
|  | Ambient Temperature | -0.43 | 0.02 | 382.83 | <0.001 |
| Time in Warm (day) | Treatment (P) | 0.18 | 0.23 | 0.39 | 0.42 |
|  | Clutch Size | 0.15 | 0.11 | 0.26 | 0.17 |
|  | Clutch Initiation Date | 0.23 | 0.12 | 0.45 | 0.06 |
|  | Embryo Age | 0.14 | 0.07 | 13.68 | 0.06 |
|  | Ambient Temperature | 0.53 | 0.04 | 224.86 | <0.001 |
| Time in Hot (day) | Treatment (P) | 1.24 | 1.8 | 0.95 | 0.49 |
|  | Clutch Initiation Date | -0.34 | 0.93 | 0.15 | 0.71 |
|  | Embryo Age | 0.40 | 0.89 | 0.22 | 0.65 |
|  | Ambient Temperature | 9.63 | 1.21 | 37.90 | <0.001 |
| Time in Cold (night) | Treatment (P) | -0.91 | 0.71 | 0.30 | 0.20 |
|  | Clutch Initiation Date | -0.34 | 0.41 | 1.18 | 0.40 |
|  | Embryo Age | 0.06 | 0.41 | 0.18 | 0.88 |
|  | Ambient Temperature | -1.03 | 0.36 | 2.69 | 0.004 |
| Time in Lukewarm (night) | Treatment (P) | -0.14 | 0.49 | 0.007 | 0.77 |
|  | Clutch Initiation Date | -0.11 | 0.26 | 1.94 | 0.67 |
|  | Embryo Age | 0.07 | 0.06 | 0.94 | 0.26 |
|  | Ambient Temperature | -0.39 | 0.07 | 34.74 | <0.001 |
| Time in Warm (night) | Treatment (P) | 2.08 | 0.82 | 2.93 | 0.01 |
|  | Clutch Size | 0.51 | 0.44 | 0.19 | 0.25 |
|  | Clutch Initiation Date | 1.41 | 0.45 | 15.19 | 0.001 |
|  | Embryo Age | -0.23 | 0.07 | 0.06 | 0.002 |
|  | Ambient Temperature | 0.81 | 0.08 | 109.51 | <0.001 |

**Figure S2.**
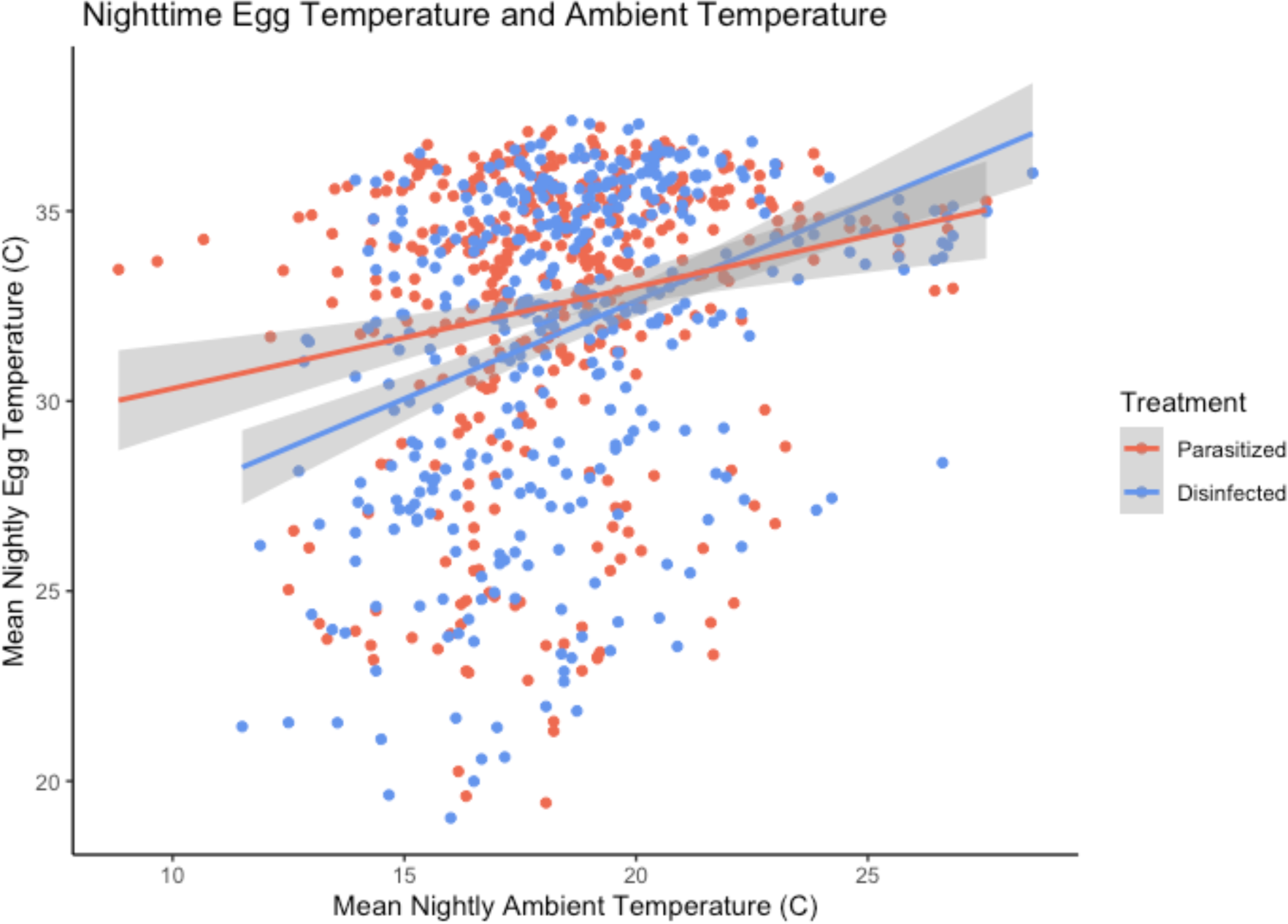
Relationship between nighttime mean egg temperature, measured with omegga loggers, and ambient temperature near the nest, measured with ibuttons, separated by parasite treatment group. Egg temperature was largely driven by ambient temperature. Figure and trend lines were made with raw data, but analyses presented in the main text include additional fixed effects and embryo age nested within nest and site as random effects to account for repeated measures within the same nest and multiple nests from the same breeding colonies.

**Figure S3.**
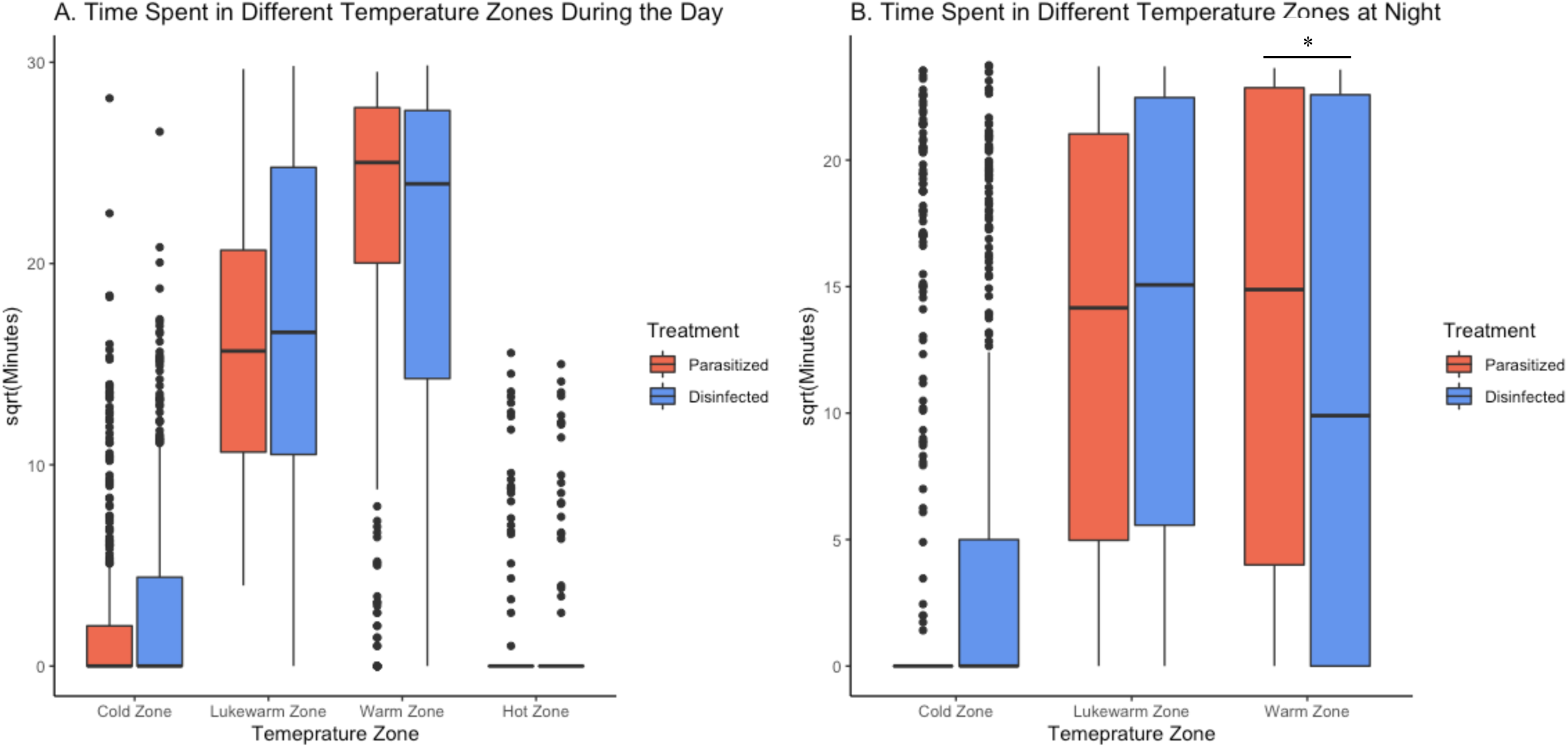
Number of minutes that eggs spent in different temperature zones, measured using omegga loggers, during A) the day and B) the night. Temperature zones are: cold (<26°C), lukewarm (26-34°C), warm (34-40.5°C), and hot (>40.5°C), representing biologically meaningful ranges during embryonic development. There were no readings in the hot zone at night. Number of minutes has been square root transformed for ease of visualization. Time spent in these zones was largely driven by ambient temperature. Parasitized nests did spend significantly more time in the warm zone at night compared to disinfected nests (*). Figures and trend lines were made with raw data, but analyses presented in the main text include additional fixed effects and embryo age nested within nest and site as random effects to account for repeated measures within the same nest and multiple nests from the same breeding colonies.

### Incubation Behavior

**Table S3.** Full model results for measures of incubation behavior. All models contained embryo age nested within nest ID and site as random effects.

Incubation Behavior:
|  |  |  |  |  |  |
| --- | --- | --- | --- | --- | --- |
| Response | Fixed Effect | Estimate | SE | F | P |
| Number of Off Bouts<br>($R^2c = 0.82$) | Treatment (P) | 3.17 | 2.97 | 1.14 | 0.29 |
|  | Clutch Size | -2.80 | 1.51 | 3.39 | 0.07 |
|  | Clutch Initiation Date | 2.09 | 1.56 | 1.78 | 0.19 |
|  | Embryo Age | 1.45 | 0.42 | 11.55 | 0.001 |
| | Ambient Temperature | -5.22 | 0.41 | 162.40 | $<0.001$ |
| Off Bout Length<br>($R^2c = 0.67$) | Treatment (P) | 3.79 | 58.46 | 0.004 | 0.95 |
|  | Clutch Size | 23.25 | 29.87 | 0.61 | 0.44 |
|  | Clutch Initiation Date | -43.93 | 31.24 | 1.98 | 0.16 |
|  | Embryo Age | -9.48 | 13.15 | 0.52 | 0.47 |
| | Ambient Temperature | 123.55 | 12.38 | 99.64 | $<0.001$ |
| Proportion of Time on the Nest | Treatment (P) | 0.01 | 0.04 | (Z) | 0.72 |
|  | Clutch Size | 0.03 | 0.02 | (Z) | 0.14 |
|  | Clutch Initiation Date | 0.02 | 0.02 | (Z) | 0.35 |
|  | Embryo Age | -0.01 | 0.01 | (Z) | 0.15 |
| | Ambient Temperature | -0.06 | 0.01 | (Z) | $<0.001$ |

### Outcomes of Incubation

**Table S4.** Full model results for per egg probability of hatching failure and days to hatch (outcomes of incubation). All models contained embryo age nested within nest ID and site as random effects.

***Outcomes of Incubation:***
|  |  |  |  |  |  |
| --- | --- | --- | --- | --- | --- |
| **Response** | **Fixed Effect** | **Estimate** | **SE** | **F** | **P** |
| Per Egg Probability of Hatching Failure<br>($R^2c = 0.27$) | Treatment (P) | 0.75 | 0.34 | 1.09 | 0.03 |
|  | Mean Daily Egg Temperature | 0.71 | 0.23 | 13.88 | 0.002 |
|  | Number Off Bouts | 0.32 | 0.14 | 6.85 | 0.02 |
|  | Clutch Size | -0.57 | 0.14 | 17.00 | <0.001 |
| Days to Hatch | Treatment (P) | -0.01 | 0.02 | 0.004 | 0.62 |
|  | Mean Daily Egg Temperature | -0.002 | 0.01 | 1.86 | 0.88 |
|  | Number Off Bouts | 0.007 | 0.01 | 0.43 | 0.50 |
|  | Clutch Size | -0.01 | 0.01 | 0.14 | 0.29 |
|  | Clutch Initiation | -0.03 | 0.01 | 6.39 | 0.01 |

## REFERENCES

Amininasab, S.M., Kingma, S.A., Birker, M., Hildenbrandt, H. & Komdeur, J. (2016). The effect of ambient temperature, habitat quality and individual age on incubation behaviour and incubation feeding in a socially monogamous songbird. Behav. Ecol. Sociobiol., 70, 1591– 1600.

Ardia, D.R. & Clotfelter, E.D. (2007). Individual quality and age affect responses to an energetic constraint in a cavity-nesting bird. Behav. Ecol., 18, 259–266.

Ardia, D.R., Pérez, J.H., Chad, E.K., Voss, M.A. & Clotfelter, E.D. (2009). Temperature and life history: Experimental heating leads female tree swallows to modulate egg temperature and incubation behaviour. J. Anim. Ecol., 78, 4–13.

Ardia, D.R., Pérez, J.H. & Clotfelter, E.D. (2010). Experimental cooling during incubation leads to reduced innate immunity and body condition in nestling tree swallows. Proc. Biol. Sci., 277, 1881–8.

Ardia, D.R., Wasson, M.F. & Winkler, D.W. (2006). Individual quality and food availability determine yolk and egg mass and egg composition in tree swallows *Tachycineta bicolor*. J. Avian Biol., 37, 252–259.

Banbura, J. & Zielinski, P. (1995). The onset of incubation and hatching asynchrony in the barn swallow *Hirundo rustica*. Ornis Fenn., 72, 174–176.

Bartroń, K. (2017). MuMIn: Multi-model inference. R package.

Bates, D., Maechler, M., Bolker, B. & Walker, S. (2015). Fitting linear mixed-effects models using lme4. J. Stat. Softw., 67, 1–48.

Bogdanova, M.I., Nager, R.G. & Monaghan, P. (2007). Age of the incubating parents affects nestling survival: An experimental study of the herring gull *Larus argentatus*. J. Avian Biol., 38, 83–93.

Boulton, R.L. & Cassey, P. (2012). How avian incubation behaviour influences egg surface temperatures: Relationships with egg position, development and clutch size. J. Avian Biol., 43, 289–296.

Bouslama, Z., Lambrechts, M.M., Ziane, N., Djenidi, R. & Chabi, Y. (2002). The effect of nest ectoparasites on parental provisioning in a north-African population of the Blue Tit *Parus caeruleus*. Ibis, 144, E73–E78.

Brommer, J.E.J., Pitala, N., Siitari, H., Kluen, E. & Gustafsson, L. (2011). Body size and immune defense of nestling Blue Tits (*Cyanistes caeruleus*) in response to manipulation of rctoparasites and food supply. Auk, 128, 556–563.

Brooks, M.E., Kristensen, K., van Benthem, K.J., Magnusson, A., Berg, C.W., Nielsen, A., et al. (2017). glmmTMB balances speed and flexibility among packages for zero-inflated generalized linear mixed modeling. R J., 9, 378–400.

Brown, C.R. & Brown, M.B. (1999). Barn Swallow (*Hirundo rustica*). In: The Birds of North America. Cornell Lab of Ornithology, Ithaca, p. 32.

Bryan, S.M. & Bryant, D.M. (1999). Heating nest-boxes reveals an energetic constraint on incubation behaviour in great tits, Parus major. Proc. R. Soc. B Biol. Sci., 266, 157.

Cantarero, A., López-Arrabé, J., Redondo, A.J. & Moreno, J. (2013). Behavioural responses to ectoparasites in pied flycatchers *Ficedula hypoleuca*: an experimental study. J. Avian Biol., 44, 1–9.

Christensen, R.H.B. (2019). Ordinal: Regression models for ordinal data. R package.

Clayton, D.H. & Tompkins, D.M. (1995). Comparative effects of mites and lice on the reproductive success of rock doves (*Columba livia*). Parasitology, 110, 195–206.

Cones, A.G., Liebl, A.L., Houslay, T.M. & Russell, A.F. (2020). Temperature-mediated plasticity in incubation schedules is unlikely to evolve to buffer embryos from climatic challenges in a seasonal songbird. J. Evol. Biol., 1–12.

Conway, C.J. & Martin, T.E. (1999). Effects of ambient temperature on avian incubation behavior, 11, 178–188.

Conway, C.J. & Martin, T.E. (2000). Evolution of passerine incubation behavior: Influence of food, temperature, and nest predation. Evolution (N. Y)., 54, 670–685.

Cooper, C.B., Mills, H., Cooper, C.B. & Mills, H. (2005). New software for quantifying incubation behavior from time-series recordings, 76, 352–356.

Cooper, C.B. & Voss, M.A. (2013). Avian incubation patterns reflect temporal changes in developing clutches. PLoS One, 8, 1–6.

De Coster, G., De Neve, L., Martín-Gálvez, D., Therry, L. & Lens, L. (2010). Variation in innate immunity in relation to ectoparasite load, age and season: a field experiment in great tits (*Parus major*). J. Exp. Biol., 213, 3012–8.

Deeming, C. (2002). Avian incubation: behaviour, environment and evolution. Oxford University Press.

Dobbs, R.C., Styrsky, J.D. & Thompson, C.F. (2006). Clutch size and the costs of incubation in the house wren. Behav. Ecol., 17, 849–856.

Douma, J.C. & Weedon, J.T. (2019). Analysing continuous proportions in ecology and evolution: A practical introduction to beta and Dirichlet regression. Methods Ecol. Evol., 10, 1412–1430.

Dube, W.C., Hund, A.K., Turbek, S.P. & Safran, R.J. (2018). Microclimate and host body condition influence mite population growth in a wild bird-ectoparasite system. Int. J. Parasitol. Parasites Wildl., 7, 301–308.

Duckworth, R.A., Belloni, V. & Anderson, S.R. (2015). Cycles of species replacement emerge from locally induced maternal effects on offspring behavior in a passerine bird. Science, 347, 875–877.

Duffey, D.C. (1983). The ecology of tick parasitism on densely nesting Peruvian seabirds. Ecology, 64, 110–119.

DuRant, S.E., Hopkins, W. a., Hepp, G.R. & Walters, J.R. (2013). Ecological, evolutionary, and conservation implications of incubation temperature-dependent phenotypes in birds. *Biol. Rev.*, Khubbard, 499–509.

Engstrand, S.M. & Bryant, D.M. (2002). A trade-off between clutch size and incubation efficiency in the Barn Swallow *Hirundo rustica*. Funct. Ecol., 16, 782–791.

Fitze, P.S., Tschirren, B. & Richner, H. (2004). Life history and fitness consequences of ectoparasites. J. Anim. Ecol., 73, 216–226.

Gabrielsen, G. & Steen, J.B. (1979). Tachycardia during egg-hypothermia in incubating ptarmigan (*Lagopus lagopus*). Acta Physiol. Scand., 107, 273–277.

Gorman, H.E., Orr, K.J., Adam, A. & Nager, R.G. (2005). Effects of incubation conditions and offspring sex on embryonic development and survival in the Zebra Finch (*Taeniopygia guttata*). Auk, 122, 1239–1248.

Haftorn, S., Reinertsen, R.E., Ornis, S., Scandinavian, S. & Apr, N. (1982). Regulation of body temperature and heat transfer to eggs during incubation. Oikos, 13, 1–10.

Hanssen, S.A., Hasselquist, D., Folstad, I. & Erikstad, K.E. (2005). Cost of reproduction in a long-lived bird: incubation effort reduces immune function and future reproduction. Proc. R. Soc. B Biol. Sci., 272, 1039–1046.

Hepp, G.R., Durant, S.E. & Hopkins, W. a. (2015). Influence of incubation temperature on offspring phenotype and fitness in birds. In: Nests, eggs, and incubation: new ideas about avian reproduction. (eds. Deeming, D. & Reynolds, S.J.). Oxford University Press, Oxford, pp. 171–178.

Hund, A.K., Aberle, M.A. & Safran, R.J. (2015a). Parents respond in sex-specific and dynamic ways to nestling ectoparasites. Anim. Behav., 110, 187–196.

Hund, A.K., Blair, J.T. & Hund, F.W. (2015b). A review of available methods and description of a new method for eliminating ectoparasites from bird nests. J. F. Ornithol., 86, 191–204.

Hund, A.K., Hubbard, J.K., Krausová, S., Munclinger, P. & Safran, R.J. (2021). Different underlying mechanisms drive associations between multiple parasites and the same sexual signal. Anim. Behav., 172, 183–196.

Killpack, T.L., Oguchi, Y. & Karasov, W.H. (2013). Ontogenetic patterns of constitutive immune parameters in altricial house sparrows. J. Avian Biol., 44, 513–520.

Lehmann, T. (1993). Ectoparasites: direct impact on host fitness. Parasitol. Today, 9, 8–13.

Levin, I.I., Fosdick, B.K., Tsunekage, T., Aberle, M.A., Bergeon Burns, C.M., Hund, A.K., et al. (2018). Experimental manipulation of a signal trait reveals complex phenotype-behaviour coordination. Sci. Rep., 8, 15533.

Liu, J., Yan, X., Li, Q., Wang, G., Liu, H., Wang, J., et al. (2013). Thermal manipulation during the middle incubation stage has a repressive effect on the immune organ development of Peking ducklings. J. Therm. Biol., 38, 520–523.

Londoño, G.A., Levey, D.J. & Robinson, S.K. (2008). Effects of temperature and food on incubation behaviour of the northern mockingbird, Mimus polyglottos. Anim. Behav., 76, 669–677.

López-Arrabé, J., Cantarero, A., Pérez-Rodríguez, L., Palma, A., Alonso-Alvarez, C., González-Braojos, S., et al. (2015). Nest-dwelling ectoparasites reduce antioxidant defences in females and nestlings of a passerine: a field experiment. Oecologia, 179, 29–41.

Loye, J.E. & Carroll, S.P. (1998). Ectoparasite behavior and its effects on avian nest site selection. Ann. Entomol. Soc. Am., 91, 159–163.

Macdonald, E.C., Camfield, A.F., Jankowski, J.E. & Martin, K. (2013). Extended incubation recesses by alpine-breeding Horned Larks: a strategy for dealing with inclement weather? J. F. Ornithol., 84, 58–68.

Mariette, M.M. & Buchanan, K.L. (2016). Prenatal acoustic communication programs offspring for high posthatching temperatures in a songbird. Science (80-.)., 353, 812–814.

Martin, T.E. (2002). A new view of avian life-history evolution tested on an incubation paradox. Proc. R. Soc. B Biol. Sci., 269, 309–316.

Martin, T.E., Auer, S.K., Bassar, R.D., Niklison, A.M. & Lloyd, P. (2007). Geographic variation in avian incubation periods and parental influences on embryonic temperature. Evolution (N. Y)., 61, 2558–2569.

Mazgajski, T. (2007). Effect of old nest material on nest site selection and breeding parameters in secondary hole nesters — a review. Acta Ornithol., 42, 1–14.

Mcculloch, J.B. & Owen, J.P. (2012). Arrhenotoky and oedipal mating in the northern fowl mite (Ornithonyssus sylviarum) (Acari: Gamasida: Macronyssidae). Parasites and Vectors, 5, 281.

Merrill, L., Ospina, E.A., Santymire, R.M. & Benson, T.J. (2020). Egg incubation temperature affects development of innate immune function in nestling American Robins (*Turdus migratorius*). Physiol. Biochem. Zool., 93, 1–12.

Møller, A.P. (1990). Effects of parasitism by a haematophagous mite on reproduction in the barn swallow. Ecology, 71, 2345–2357.

Møller, A.P. (1993). Ectoparasites increase the cost of reproduction in their hosts. J. Anim. Ecol., 62, 309–322.

Morris, B.M., Tollerud, E., Sipőcz, B., Deil, C., Douglas, S.T., Medina, J.B., Vyhmeister, K, et al. (2018). Astroplan: an open source observation planning package in Python.” Astron J., 155, 3, 128.

Morrison, E.S., Ardia, D.R. & Clotfelter, E.D. (2009). Cross-fostering reveals sources of variation in innate immunity and hematocrit in nestling tree swallows *Tachycineta bicolor*. J. Avian Biol., 40, 573–578.

Nilsson, J.Å.J. (2003). Ectoparasitism in marsh tits: costs and functional explanations. Behav. Ecol., 14, 175–181.

Nord, A. & Nilsson, J. Åke. (2012). Context-dependent costs of incubation in the pied flycatcher. Anim. Behav., 84, 427–436.

Nord, A. & Williams, J.B. (2015). The energetic costs of incubation. In: Nests, Eggs, and Incubation: New Ideas about Avian Reproduction (eds. Deeming, C. & Reynolds, S.J.). Oxford University Press, Oxford, pp. 152–170.

Olson, C.R., Vleck, C.M. & Vleck, D. (2006). Periodic cooling of bird eggs reduces embryonic growth efficiency. Physiol. Biochem. Zool., 79, 927–936.

Oppliger, A., Richner, H. & Christe, P. (1994). Effect of an ectoparasite on lay date, nest-site choice, desertion, and hatching success in the great tit (*Pants major*). Behav. Ecol., 5, 130– 134.

Owen, J.P., Nelson, A.C. & Clayton, D.H. (2010). Ecological immunology of bird-ectoparasite systems. Trends Parasitol., 26, 530–539.

Palacios, M.G., Cunnick, J.E., Vleck, D. & Vleck, C.M. (2009). Ontogeny of innate and adaptive immune defense components in free-living tree swallows, *Tachycineta bicolor*. Dev. Comp. Immunol., 33, 456–63.

Peralta-Sánchez, J.M., Martín-Platero, A.M., Wegener-Parfrey, L., Martínez-Bueno, M., Rodríguez-Ruano, S., Navas-Molina, J.A., et al. (2018). Bacterial density rather than diversity correlates with hatching success across different avian species. FEMS Microbiol. Ecol., 94, 1–13.

Pérez, J.H., Ardia, D.R., Chad, E.K. & Clotfelter, E.D. (2008). Experimental heating reveals nest temperature affects nestling condition in tree swallows (*Tachycineta bicolor*). Biol. Lett., 4, 468–471.

Proctor, H. & Owens, I. (2000). Mites and birds: diversity, parasitism and coevolution. Trends Ecol. Evol., 15, 358–364.

Pryor, L.J.E. & Casto, J.M. (2017). Ectoparasites as developmental stressors: Effects on somatic and physiological development. J. Exp. Zool. Part A Ecol. Integr. Physiol., 327, 311–321.

Reid, J.M., Monaghan, P. & Ruxton, G.D. (2000). Resource allocation between reproductive phases: the importance of thermal conditions in determining the cost of incubation. Proc. R. Soc. B, 267, 37–41.

Reid, J.M., Ruxton, G.D., Monaghan, P. & Hilton, G.M. (2002). Energetic consequences of clutch temperature and clutch size for a uniparental intermittent incubator: The starling. Auk, 119, 54–61.

Richner, H. & Heeb, P. (1995). Are clutch and brood size patterns in birds shaped by ectoparasites? Oikos, 73, 435.

Ruiz-Castellano, C., Tomás, G., Ruiz-Rodríguez, M., Martín-Gálvez, D. & Soler, J.J. (2016). Nest material shapes eggs bacterial environment. PLoS One, 11, 1–21.

Skutch, A.F. (1962). The constancy of incubation. Wilson Bull., 74, 115–152.

Smith, H.G. & Montgomerie, R. (1992). Male incubation in barn swallows: The influence of nest temperature and sexual selection. Condor, 94, 750–759.

Stambaugh, T., Houdek, B.J., Lombardo, M.P., Thorpe, P.A. & Caldwell Hahn, D. (2011). Innate immune response development in nestling tree swallows. Wilson J. Ornithol., 123, 779–787.

Szabó, K., Szalmás, A., Liker, A. & Barta, Z. (2002). Effects of haematophagous mites on nestling house sparrows (*Passer domesticus*). Acta Parasitol., 47, 318–322.

Tøien, Ø. (1993). Control of shivering and heart rate in incubating bantam hens upon sudden exposure to cold eggs. Acta Physiol. Scand., 149, 205–214.

Tomás, G., Martín-Gálvez, D., Ruiz-Castellano, C., Ruiz-Rodríguez, M., Peralta-Sánchez, J.M., Martín-Vivaldi, M., et al. (2018). Ectoparasite activity during incubation increases microbial growth on avian eggs. Microb. Ecol., 76, 555–564.

Tripet, F., Glaser, M. & Richner, H. (2002). Behavioural responses to ectoparasites: time-budget adjustments and what matters to Blue Tits *Parus caeruleus* infested by fleas. Ibis (Lond. 1859)., 144, 461–469.

Tripet, F., Richner, H. & Tripet, F. (1997). Host responses to ectoparasites: Food compensation by parent Blue Tits. Oikos, 78, 557.

Turbek, A.S.P., Hund, A.K., Hernandez, M., Hubbard, J.K. & Hubbard, J.K. (2019). The double incubation of separate nests, 182.

Turner, J.S. (1997). On the thermal capacity of a bird’s egg warmed by a brood patch. Physiol. Zool., 70, 470–480.

Visser, M.E. & Lessells, C.M. (2001). The costs of egg production and incubation in great tits (*Parus major*). Proc. R. Soc. B Biol. Sci., 268, 1271–1277.

Voss, M.A., Reed Hainsworth, F. & Ellis-Felege, S.N. (2006). Use of a new model to quantify compromises between embryo development and parental self-maintenance in three species of intermittently incubating passerines. J. Therm. Biol., 31, 453–460.

Wada, H., Kriengwatana, B., Allen, N., Schmidt, K.L., Soma, K.K. & MacDougall-Shackleton, S.A. (2015). Transient and permanent effects of suboptimal incubation temperatures on growth, metabolic rate, immune function, and adrenocortical responses in zebra finches. J. Exp. Biol., 218, 2847–2855.

Wiebe, K., Wiehn, J. & KorpimÄki, E. (1998). The onset of incubation in birds: can females control hatching patterns? Anim. Behav., 55, 1043–52.

Williams, K., Sudnick, M., Anderson, R. & Fitschen-Brown, M. (2020). Experience counts: The role of female age in morning incubation and brooding behavior in relation to temperature. J. Avian Biol., 51, 1–12.

Zuur, A., Leno, E.N., Walker, N., Saveliev, A.A. & Smith, G.M. (2009). *Mixed effects models and extensions in ecology with R*. Springer New York, New York.

